# Brief report on differential toxicity of legacy and second-generation PFAS on dopaminergic neurons and mitochondria

**DOI:** 10.1101/2025.10.08.681169

**Authors:** Napissara Boonpraman, Da-Woon Kim, Nathan C Kuhn, Souvarish Sarkar, Shreesh Raj Sammi

## Abstract

Per/polyfluoroalkyl substances (PFAS) are anthropogenic chemicals that have shown extensive usage. Owing to widespread use and resistance to environmental degradation, they have become a hazard with respect to the environment and human health. While the legacy PFAS are being phased out, they are being replaced by second-generation PFAS that are considered safer alternatives. However, the lack of information pertaining to the underlying mechanisms for legacy and especially second-generation PFAS exacerbates the risk. This study investigates legacy and second-generation PFAS to determine their individual effects on neurotoxicity and mitochondrial respiration. Legacy PFAS, perfluorooctanoic acid (PFOA), perfluorononanoic acid (PFNA), perfluorodecanoic acid (PFDA), and second-generation PFAS, GenX, and ADONA were employed, and studies were conducted using *Caenorhabditis elegans* and mitochondria isolated from Rat brain for Complex I to IV. It was found that legacy PFAS, PFNA, and PFDA were neurotoxic, with neurotoxicity proportional to chain length. PFOA, PFNA, and PFDA also exhibited significant inhibition to most of the mitochondrial complexes. Whereas in the case of second-generation PFAS ADONA, and GenX, it was only limited to the inhibition of Complex IV. GenX only exhibited neurodegeneration at very high doses. Our findings conclude that the legacy and second-generation PFAS might have significant neurotoxic implications. While the targets are similar to some extent, the mechanisms between the two classes are distinct. Our study the sets the stage to evaluate further the combined effect of PFAS (legacy and emergent) to fill in informational gaps pertaining to their safety.

## Introduction

Per/polyfluoroalkyl substances (PFAS) are man-made chemicals^1, 2^. Their characterstics, such as resistance to degradation^3, 4^ and the ability to repel oil and water^5, 6^, contribute to their widespread use in industrial and consumer applications^7, 8^. However, due to their unrestricted use, PFASs have become a major environmental contaminant and a risk to human and ecological health. A 2007 study employing 2094 serum samples showed the presence of PFAS in over 98% of samples^9^. Legacy PFAS such as perfluorooctanoic (PFOA)^10^ acid and perfluoro sulfonic acid (PFOS)^11^ were voluntarily phased out in 2006 and 2002, respectively. However, they were soon replaced by alternative PFAS. Before transitioning to the second-generation PFAS, shorter-chain PFAS such as perfluorobutanoic acid (PFBA) were introduced as PFAS replacements^12^. Soon, these were replaced by second-generation PFAS such as GenX (hexafluoropropylene oxide dimer acid)^13-15^ and ADONA (4,8-dioxa-3H-perfluorononanoic acid)^14, 15^. These newer PFAS are often short-chain or branched and are considered less toxic than their predecessors. However, several studies have questioned the safety of second-generation PFAS, and some have suggested that it may be a regrettable substitution^16-18^. Predominant legacy PFAS are typically linear and have longer chain lengths. They have been classified based on functional groups, such as PFSA (perfluoroalkyl sulfonic acid) and PFCA (perfluoroalkyl carboxylic acid). Before transitioning to the second-generation PFAS, shorter-chain PFAS such as perfluorobutanoic acid (PFBS) were introduced as PFAS replacements.

Legacy PFAS have an immense half-life in the environment, ensuring their presence as part of exposomes for the decades to come^19-21^. Further complicating the situation, they are accompanied by the newer, second-generation PFAS that have not been evaluated for their toxicity and adverse outcomes. Hence, it is essential to assess the legacy and newer PFAS for toxicity and to determine any common mechanisms. This brief study evaluates the differential toxicity of legacy and newer PFAS for their effect on mitochondrial respiration and dopaminergic neurons, with a focus on elucidating the toxicity and commonalities between the mechanisms.

## Material and Methods

### Culture and maintenance of Strains

*Caenorhabditis elegans* strains, BZ555 (egls1[dat-1p::GFP]), and *Escherichia coli* OP50, were procured from the Caenorhabditis Genetics Centre (University of Minnesota, Minnesota), grown on Nematode growth medium (NGM), and cultured at 22°C. A synchronized population of worms was obtained by sodium hypochlorite treatment. Embryos were incubated overnight at 15°C in M9 buffer to obtain L1 worms.

### Neurodegeneration Assay

L1-stage worms were treated with different concentrations of PFAS in K medium complete as described previously^22-25^. Three perfluoroalkyl carboxylic acids PFAS, perfluorooctanoic acid (PFOA), perfluorononanoic acid (PFNA), and perfluorodecanoic acid (PFDA), along with two second-generation PFAS, 4,8-dioxa-3H-perfluorononanoic acid (ADONA) and hexafluoropropylene oxide dimer acid (GenX), were tested. 10,000-ppm stock of PFAS was prepared in methanol and diluted further to doses ranging from 25 to 75 ppm. For extended-range studies (25 to 400 ppm), ADONA and GenX stocks were prepared at a concentration of 40,000 ppm. Approximately 175 L-1 worms were treated for 72 hours in 12-well plates, with 500 μL per well. The concentration of methanol was adjusted accordingly for each experimental group, with the final concentration being 1%. Neurodegeneration was scored as described previously^23-25^. Briefly, the treated worms were washed three times using M9 buffer and anesthetized by adding 7 μL of 100 mM sodium azide in 100 μL of worm suspension. Counting of neurons was done for all neuron types i.e., Cephalic sensilla (CEP), Anterior deirid (ADE), and posterior deirid (PDE), using a FITC filter. *C. elegans* have eight DA neurons, 2 ADE,, 4 CEP and 2 PDE ^26^. The percentage of intact neurons (PIN) was calculated for a minimum of 20 worms per group (independent biological replicates).

### Assay for mitochondrial enzyme activity

Assessment of mitochondrial complex activity for complexes I to IV was done as described by Spinazzi et al., 2012^27^. Briefly, mitochondria were isolated from rat brain and treated with different doses of PFAS for 15 minutes with doses 25, 75 ppm. The substrate was added, and optical density was read every 15 seconds for 3 minutes. Relative enzyme activity was calculated as: (Δ Absorbance/min × 1,000)/[(extinction coefficient × volume of sample used in ml) × (sample protein concentration in mg/ml)]. Graphs were plotted as relative enzyme activity, normalized with respect to the control.

## Results

### Legacy PFAS with more than eight carbon atoms exhibit dopaminergic neurotoxicity

In view of the previous studies indicating dopaminergic cell loss upon exposure to perfluorooctanesulfonate^24^, we compared the legacy and second-generation PFAS for their effect on the dopaminergic cell loss. Three legacy perfluoroalkyl carboxylic acids were tested along with two second-generation PFAS, ADONA and GenX. Amongst tested PFAS, only PFNA and PFDA exhibited neurodegeneration. PFNA showed significant decrease in percentage of intact neurons (PIN) at doses 50 and 75 ppm (Figure 1 C). PFDA exhibited a relatively severe effect on dopaminergic neurons, exhibiting a significant decrease in PIN at doses 25, 50 and 75 ppm (Figure 1D). These results indicated that PFAS neurotoxicity is proportional to the chain length.

**Figure 1:**
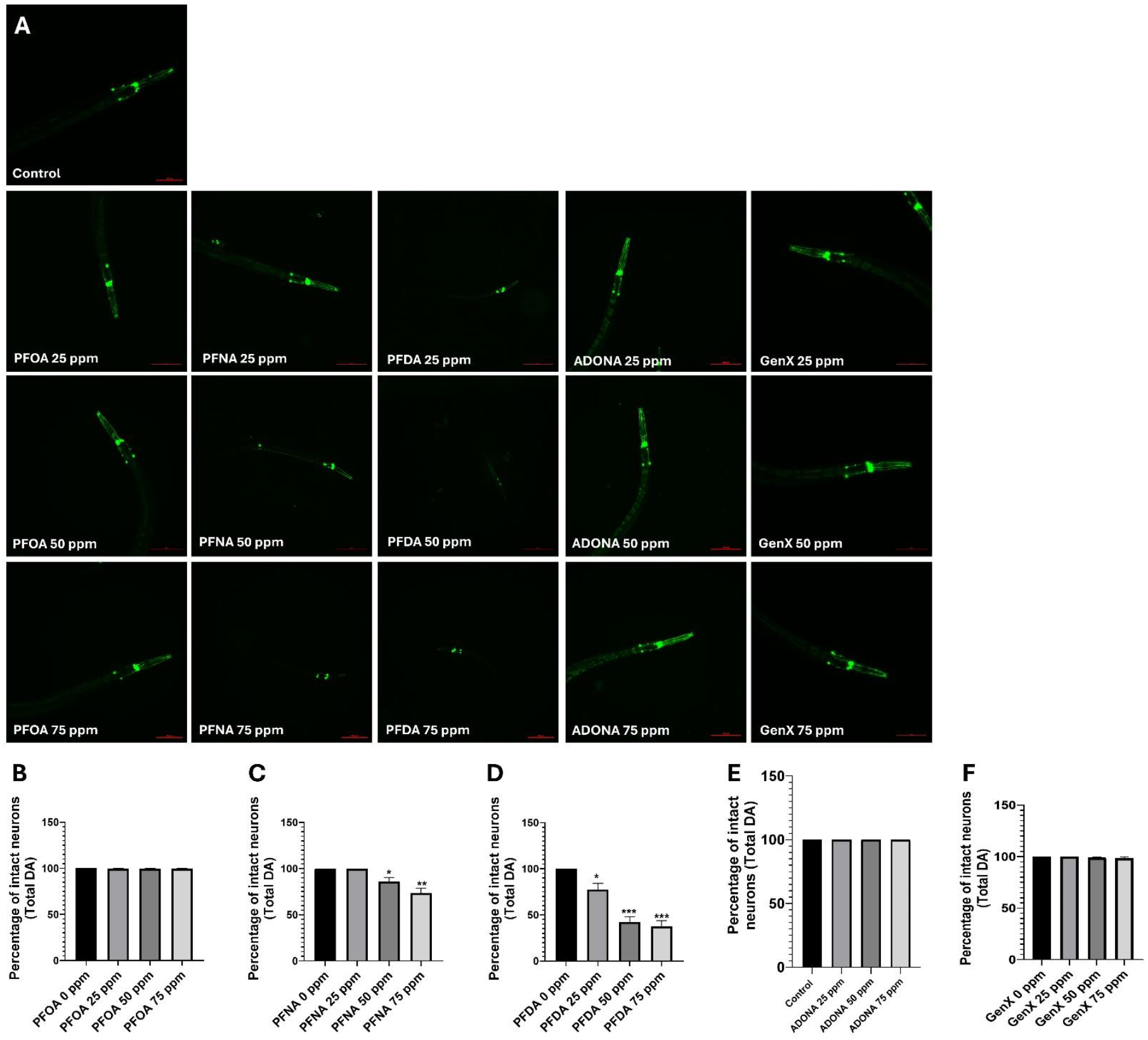
Legacy PFAS greater than eight carbon atoms exhibit dose-dependent neurotoxicity in dopaminergic neurons. C. elegans strain BZ555 were treated with three legacy and two second-generation PFAS to test for the effect on dopaminergic cell loss (representative micrographs (A)). PFAS above eight carbon, PFNA (C) and PFDA (D) exhibited exposure-dependent neurotoxicity that exhibited an upward trend with an increase in carbon number, whereas eight-carbon PFAS, PFOA (B), and second-generation PFAS, ADONA (E) and GenX (F) were devoid of any neurotoxicity within the exposure range, 25 to 75 ppm. L1 worms were subjected to PFOS for 72 h. Data presented as mean + S.E.M. Data was analyzed using one-way ANOVA followed by Dunnett’s post hoc test. *p<.05, **p<.005, and ***p<.001 (n =3). Scale bar represents 100 μM.

### GenX exhibits dopaminergic cell loss at very high concentrations

After testing the effect of PFAS at concentration range, 25 to 75 ppm, we found that the second-generation PFAS have no effect on dopaminergic cells. Next we questioned, if higher concentrations might afflict dopaminergic cells. Hence, we tested the neurotoxicity of ADONA and GenX up to 400 ppm concentrations. We found that while ADONA (Figure 1 B) did not show any dopaminergic lesions, GenX did show slight neurodegeneration at 400 ppm (Figure 2 C).

**Figure 2:**
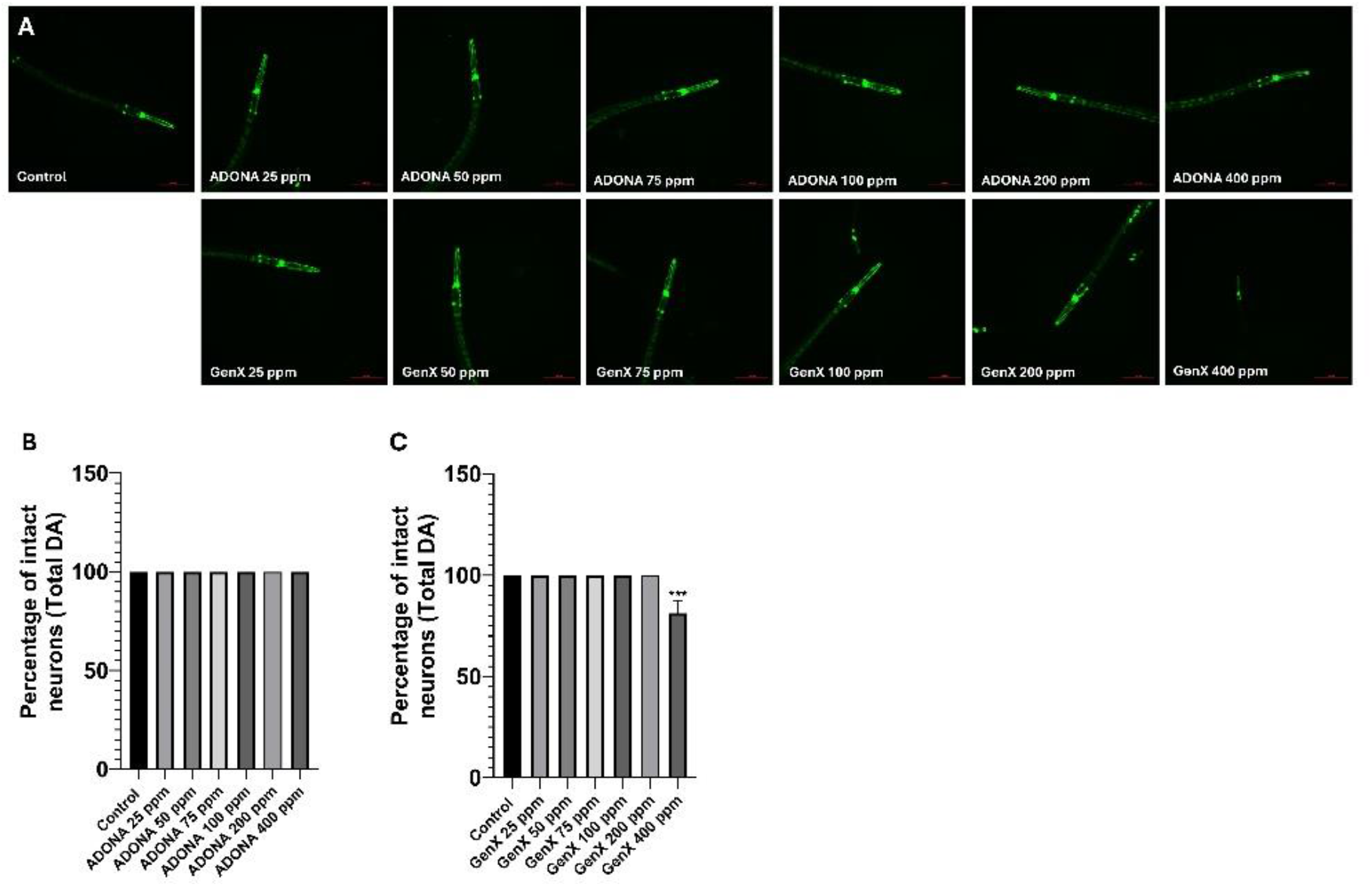
Second-generation PFAS exhibit no to a little neurotoxicity over extended range. C. elegans strain BZ555 were treated with second-generation PFAS, ADONA, and GenX to test for the effect on dopaminergic cell loss (representative micrographs (A)). While ADONA was devoid of any neurotoxicity, even at 400 ppm (B), GenX did exhibit some neurotoxicity at the highest dose of 400 ppm (C). L1 worms were subjected to PFOS for 72 h. Data presented as mean + S.E.M. Data was analyzed using one-way ANOVA followed by Dunnett’s post hoc test. ***p<.001 (n =3). Scale bar represents 100 μM.

### Effect of PFAS on mitochondrial enzyme activity

Dopaminergic cells have a high energy demand^28^ and the affliction of mitochondria or mitochondrial enzyme activity has been associated with dopaminergic cell loss and Parkinson’s disease, which is manifested with dopaminergic cell loss^29-31^. Hence, we tested the effect of PFAS on mitochondrial enzyme activity by acutely exposing mitochondria isolated from the rat brain to 25 and 75 ppm. PFOA exhibited significant dose dependent inhibition of mitochondrial complex III and IV in an exposure-dependent manner (Figure 3 A). PFNA showed inhibition of mitochondrial complex I, II, III, and IV (Figure 3 B). PFDA exhibited inhibition of complexes II, III, and IV (Figure 3 C). ADONA treatment was only limited to inhibition of complex IV (Figure 3 D). Similarly, GenX also exhibited inhibition of Complex IV only (Figure 3 E), which was less severe in comparison to ADONA and tested legacy PFAS.

**Figure 3:**
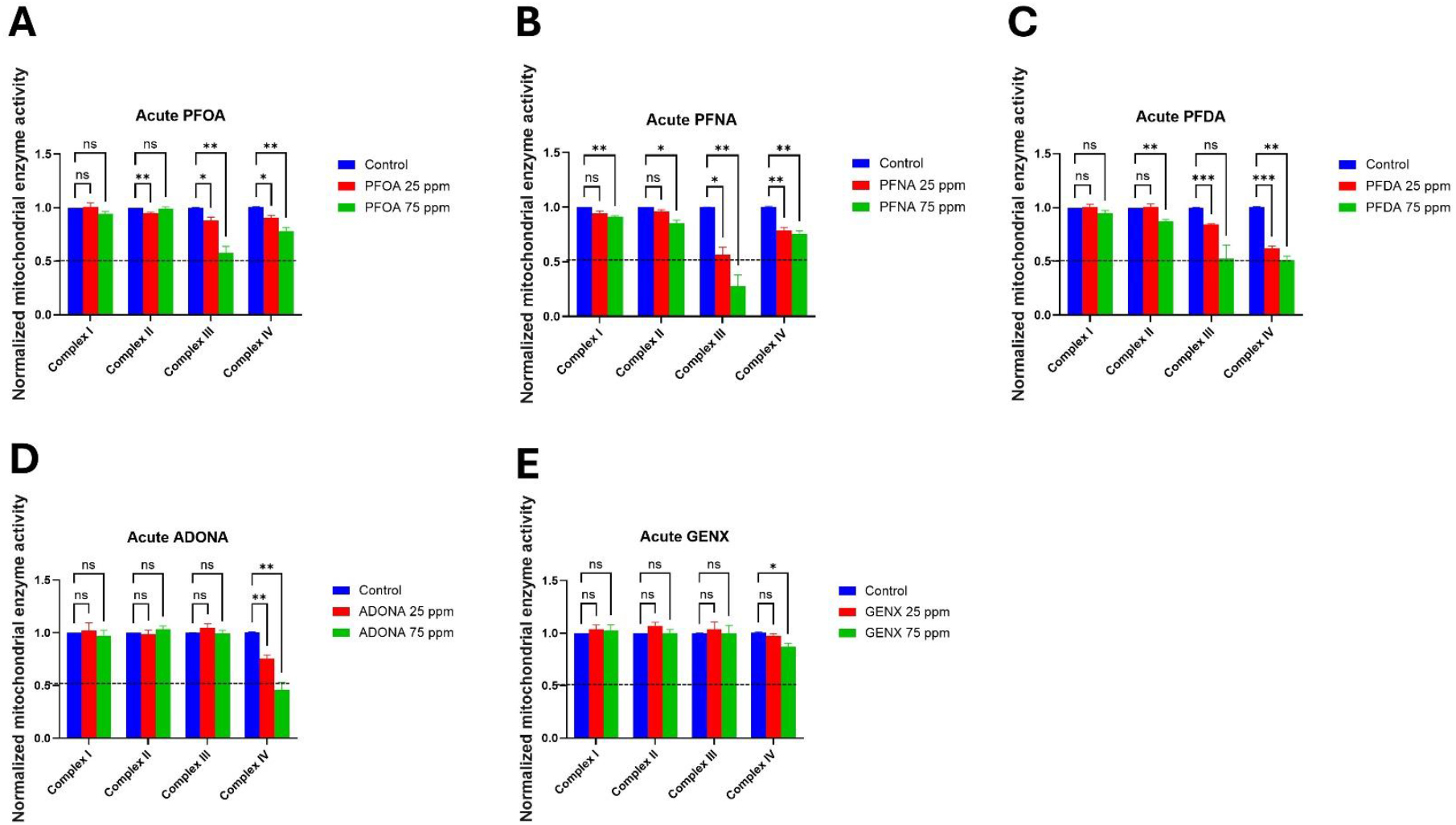
Effect of Legacy and second-generation PFAS on mitochondrial enzyme activity. Mitochondria isolated from rat brain were acutely subjected to legacy and second-generation PFAS (25, 75 ppm) for 15 minutes, and enzymatic activity for mitochondrial complexes I, II, III, and IV was assessed. PFOA exhibited slight inhibition of complex II at 25 ppm, only while a exposure-dependent increase in enzymatic inhibition was observed for complex III and IV (A). The extent of mitochondrial enzyme affliction was even higher in case of PFNA, exhibiting inhibition of complex I and II at 75 ppm only, whereas exposure-dependent increased inhibition was observed for complex III and IV, which was higher than PFOA (B). PFDA exhibited enzyme inhibition in case of higher dose (75 ppm) of complex II. While inhibition of complex III was also observed at 25 and 75 ppm, the effect was statistically insignificant at 75 ppm. Complex IV was found to be inhibited in an exposure-dependent manner at both 25 and 75 ppm (C). ADONA did not exhibit any effect on complexes I, II, and III, whereas it had a significant effect on complex IV in an exposure-dependent manner (D). GenX exposure did not exhibit any effect on complexes I, II, and III, whereas a significant effect on complex IV was observed only at 75 ppm (E). Data presented as mean + S.E.M. Data were analyzed using one-way ANOVA followed by Dunnett’s post hoc test. *p<.05, **p<.005, and ***p<.001 (n =3). Scale bar represents 100 μM.

## Discussion

First synthesized in 1930s^32^, PFAS though tremendously useful in industrial and consumer scenarios has become a huge environmental and health concern^33, 34^. While legacy PFAS have been discontinued, they are being replaced by the emergent or second-generation PFAS such as ADONA and GenX, which are typically short-chain and branched. This study tests the effect of legacy and second-generation PFAS for mechanistic commonalities pertaining to dopaminergic cell loss and mitochondrial dysfunction. Earlier studies on a prominent legacy PFAS, perfluoro-octanesulfonate have shown dopaminergic neurotoxicity and mitochondrial affliction^24^.

We questioned if there are any mechanistic similarities amongst legacy PFAS and how does it translate to the second-generation PFAS, which are deemed to be less toxic. To begin with, we ascertained the effect of legacy perfluoroalkyl carboxylic acids, PFOA, PFNA, and PFDA and ADONA, and GenX on dopaminergic neurons in *C. elegans* within the dose range. 25 to 75 ppm. Amongst the tested PFAS, only PFNA, and PFDA exhibited dopaminergic cell loss. The severity of neurotoxicity was higher in PFDA, in comparison to the PFNA. This is in line with the previous studies confirming similar trend, that short-chain PFAS are relatively less toxic^35^. Given that second-generation PFAS were devoid of any neurotoxicity in this dose range, we next tested the effect of these PFAS at very high exposures, where we observed that GenX alone afflicted dopaminergic cells at 400 ppm. While these results show that short-chain PFAS are less toxic, it is actually concerning due to the fact that short-chain PFAS can achieve up to 88.8 % concentration in water^36^. This is owing to their greater solubility in comparison to the long-chain PFAS, due to smaller chain length, which curtails hydrophobicity^37^. While the addition of an oxygen atom in ADONA and GenX is perceived as a vulnerability to allow breakdown into smaller fragments^38^; inadvertently it is likely to enhance solubility in the aqueous phase^39^. Next, we tested the effect of PFAS on mitochondrial enzyme activity. The affliction of mitochondria or mitochondrial function has been attributed to neurodegenerative disorders^40-42^, such as Parkinson’s disease^43-45^. Hence, we tested the effect of acute exposure on mitochondrial enzyme activity to see if there is a correlation with dopaminergic neurotoxicity. Our studies did show that the PFAS exhibited curtailed enzyme activity, mostly for complex III and IV. Legacy PFAS, PFNA and PFDA exhibited enzyme inhibition of mitochondrial complex II, III, and IV. Notably, these were the PFAS that showed significant neurodegeneration. Our studies showed that the mechanisms of toxicity amongst legacy and second-generation PFAS are distinct. While legacy PFAS seem to have some correlation between mitochondrial enzyme inhibition and neurodegeneration, suggesting a plausible factor, second-generation completely circumvented this trend. Slightly neurotoxic GenX (at high concentrations) was least inhibitive to mitochondrial enzyme, where as ADONA that was devoid of neurotoxicity, exhibited enhanced inhibition of mitochondrial complex IV. An overall comparison for the effect on mitochondrial inhibition and neurotoxicity points out that complex III might be an important factor.

Overall our findings clearly show that PFAS have an effect on the hallmarks associated with neurodegenerative disorders. Yet, deeper research is needed to particularly with regards to other key markers such as α-synuclein to shed more light on how PFAS, legacy and second-generation affect the pathology in neurodegeneration and ascertain the commonalities in underlying mechanisms.

## Acknowledgement

Nematode strains employed in this study were provided by the C. elegans Genetics Center (CGC), University of Minnesota, MN, USA, funded by the NIH National Center for Research Resources (NCRR). We also thank Michelle Gartland and Betsy Matazel for providing administrative support. This work was supported by the National Institute of Environmental Health Sciences (R00ES032488 to S.R.S).

## Notes

### Competing Interest Statement

The authors have declared no competing interest.

